# Comparative Evaluation of DDA- and DIA-Based Proteomic Workflows in Beryllium-Related Lung Disease

**DOI:** 10.64898/2026.06.04.730108

**Authors:** Danielle O Weise, Kanika Gupta, Timothy J Griffin, Pratik D Jagtap, Margaret M Mroz, Reid Wagner, Joshua D Macaluso, S Mehta, Lisa A Maier, L Li, B E Vestal, M Bhargava

## Abstract

We compared traditional data-dependent acquisition mass spectrometry (DDA-MS) with the increasingly adopted data-independent acquisition (DIA-MS) to evaluate their relative utility for large-scale quantitative biofluid proteomics of lung compartments, specifically paired bronchoalveolar lavage (BAL) cells and bronchoalveolar lavage fluid (BALF). Using beryllium-related granulomatous lung disease as a focused model, we analyzed BALF and BAL cells from beryllium-sensitized (BeS) individuals using both acquisition strategies to assess proteome depth, quantitative completeness, and analytical robustness.

In BAL cells, 5,640 proteins were identified by DDA-MS and 5,227 by DIA-MS; however, DIA-MS yielded markedly improved quantitative completeness, with 5,178 proteins (∼99%) quantified across all samples compared with 3,539 (∼63%) quantified by DDA-MS. While 3,397 proteins were quantified by both methods, DIA-MS uniquely quantified 1,781 lower-abundance proteins. Proteins identified by both DIA and DDA-MS approaches revealed pathways associated with granulomatous inflammation, including Toll-like receptor, clathrin-mediated endocytosis, sirtuin, and C-type lectin receptor signaling, whereas DIA-MS resolved additional pathways, such as the complement cascade, coagulation system, and JAK/IL–6–type cytokine signaling. In BALF, although more proteins were identified by DDA-MS than by DIA-MS (2,069 vs 1,742), DIA-MS achieved greater quantitative completeness, with 1,695 proteins quantified across all samples compared with 1,050 using DDA-MS, underscoring its suitability for biomarker-oriented analyses in lung fluid compartments.

Together, these results support DIA-MS as a robust and sensitive platform for quantitative lung proteomics and discovery of disease-relevant protein signatures.

## Introduction

Lung tissue provides the most direct insight into the biology of inflammatory and fibrotic lung diseases. However, acquisition of lung tissue typically requires invasive procedures such as surgical lung biopsy, limiting its use in routine clinical practice and large-scale studies. As a result, minimally invasive approaches are preferred, with bronchoscopy and bronchoalveolar lavage (BAL) enabling sampling of distal lung compartments. Bronchoalveolar lavage fluid (BALF) and associated cellular fractions therefore represent clinically accessible biofluid compartments that reflect local immune and inflammatory processes within the lung.

Granulomatous diseases, including those of known etiology, such as chronic beryllium disease (CBD), and those of unknown etiology, such as sarcoidosis and hypersensitivity pneumonitis (HP), affect individuals across diverse populations. These conditions share key immunopathogenic mechanisms, including a dominant Th1-polarized CD4⁺ T-cell response characterized by elevated production of IFN-γ and TNF-α, which drive granuloma formation and maintenance [1–4]. Antigen presentation via MHC class II molecules, with disease-associated HLA alleles—such as HLA-DPB1*E69 in CBD and its precursor, beryllium sensitization (BeS), and HLA-DRB1 variants in sarcoidosis—confers susceptibility [5–7]. Persistent antigen exposure results in chronic T-cell activation and eventual exhaustion, marked by upregulation of inhibitory receptors such as PD-1, contributing to immune dysregulation and disease chronicity [8, 9]. CBD is caused by occupational beryllium (Be) exposure. The precursor to CBD, BeS, progresses to CBD at an estimated rate of 6–8 % per year [10]. Studying BeS and CBD offers unique insights into immune mechanisms underlying granulomatous inflammation Because proteins orchestrate the cellular processes underlying granulomatous inflammation, comprehensive proteomic characterization is essential to elucidate these mechanisms. BALF and BAL cells provide a minimally invasive, clinically relevant biofluid window into the molecular and cellular mechanisms driving granulomatous lung inflammation. Several targeted proteomic platforms assess predefined sets of proteins using different detection methods—such as modified single-stranded DNA aptamers [11, 12], dual antibody–DNA conjugates[13], and next-generation immunoassays, such as the Nucleic-acid-Linked Immuno-Sandwich Assay (NULISA), which enables single-molecule counting with attomolar sensitivity [14]. By contrast, mass spectrometry (MS)-based proteomics does not require *a priori* candidates to test and has advanced substantially with high-resolution, faster MS instruments, more sophisticated bioinformatic algorithms, and improved automated biospecimen processing, thereby enhancing the reproducibility of MS proteomics. Specifically, tandem MS via Data-Independent Acquisition (DIA), compared with Data-Dependent Acquisition (DDA) of spectral data, offers distinct benefits and greater rigor for quantitative proteomic [15, 16]. Here, we directly compare contemporaneous DIA-MS and DDA-MS workflows using two well-established software platforms, EncyclopeDIA[16] and FragPipe [17], to evaluate their performance in the quantitative analysis of BAL cells and BALF from beryllium-exposed individuals.

## Methods

The National Jewish Health (NJH) Institutional Review Board (IRB HS-2466) approved the study protocols for biospecimen collection, and the BRNAY IRB (File #23-02-516-528) approved the current study.

### Participants and samples

We examined paired BALF and BAL cells from six subjects with beryllium sensitization (BeS), defined as evidence of confirmed abnormal blood or BAL beryllium lymphocyte proliferation test (BeLPT) results but without evidence of granulomatous inflammation on lung biopsy [18]. Of the six stored samples, two had BAL performed in 2023, two in 2012, and two in 2003. The samples were stored at -80°C without freeze-thaw cycles. In addition, we compared BAL cells collected in 2016 and preserved by cryopreservation or flash-freezing in liquid nitrogen, then stored at -80 °C, to assess the impact of preservation on protein yield.

### BALF and BAL cell processing for LC-MS/MS

BAL fluid was processed using a well-established protocol designed to enrich for medium- and low-abundance proteins, minimize protein loss, and yield high-quality trypsin-digested peptides free of contaminants [19], with minor modifications. Specifically, we used the High-Select Top 14 depletion resin instead of the Seppro IgY14 spin columns to enrich for medium- and low-abundance proteins. For BAL cells, sample preparation was performed as described previously [20], with the following modifications. Cell pellets were washed three times in phosphate-buffered saline (PBS) at 4°C and centrifuged at 500 g. The whole cell protein isolate was obtained using cell lysis buffer [7 M urea, 2 M thiourea, 0.4 Tris pH 8, 20% acetonitrile, 10 mM Tris (2-carboxyethyl) phosphine (TCEP), 40 mM chloroacetamide, and 1x HALT protease and phosphatase inhibitor (Thermo Fisher Scientific)], followed by sonication at 30% amplitude for 5 seconds, and then pressure cycled (35 kPSI for 20 seconds, 0 kPSI for 10 seconds; 60 cycles, 37°C) on a Barocycler NEP2320. After centrifugation (15,000 g, 10 min), the protein concentrations were determined using the Bradford assay. Peptides derived from 5 μg cellular proteins after overnight trypsin (Promega) digestion (enzyme-to-protein ratio of 1:40) at 37°C were acidified with 0.3% formic acid (FA), cleaned by Stage-Tip with an MCX-like SDB-RPS STAGE tip [21], and dried in a vacuum concentrator and frozen.

### Mass spectral data acquisition

We reconstituted the dried peptides in load solvent (H_2_O: acetonitrile (ACN): formic acid (FA), 97.9:2:0.1), and analyzed 200 nanograms of each sample by capillary LC-MS with a Thermo Fisher Scientific, Inc (Waltham, MA) Dionex UltiMate 3000 RSLCnano system on-line with an Orbitrap Eclipse mass spectrometer (Thermo Scientific, Waltham MA) for both DDA- and DIA-MS. All acquisition parameters were defined using established instrumental methods and applied consistently across samples within each acquisition strategy to enable direct comparison between DDA-MS and DIA-MS workflows.

a. *Orbitrap Eclipse DDA-LC-MS/MS:* We analyzed each sample using FAIMS (high-field asymmetric waveform ion mobility) in standard-resolution mode, as previously described[19], [22], with modifications. Changes included extending the column length to 45 cm (from 40 cm), reducing the gradient time to 77 min (from 122 min), lowering the profile for 21% B to 50 min (from 90 min) and 35% B to 75 min (from 120 min), decreasing the flow rate to 350 nL/min from 0-2 minutes (from 400 nL/min), changing the CV to -70 (from -75), increasing the FAIMS CV dwell time to 2 s (from 1 s), changing the MS1 profile to 400-1440 m/z (from 381-1400) and switching MS2 detection from Orbitrap to ion trap for faster scans. Additional adjustments included reducing dynamic exclusion to 30 s (from 45 s), increasing the charge states to 2-6 (from 2-5), lowering the precursor intensity threshold to 5000 counts (from 2.5 × 10^4), and adjusting AGC and injection times to balance speed and depth of coverage.
b. *Orbitrap Eclipse DIA- LC-MS/MS:* We analyzed each sample above with DIA-MS as described previously[15, 23] with modifications. Peptides were injected directly in the load solvent, and gradient separation was performed identically to the DDA studies above. Unlike the published method, which employed 10 m/z isolation windows with 30 MS2 scans per cycle, we implemented 12 Da windows with a 2 Da overlap and acquired 75 MS2 scans per cycle. Additionally, our collision energy was set to 33%, compared to 27% in their workflow. We also used a 3-second loop duration rather than a fixed 30-window loop. Finally, our AGC targets were adjusted to 100% (4 × 10^5) for MS1 and 800% for MS2, whereas Kleiner et al. used 300% for MS1 and 100% for MS2. These modifications were designed to improve sensitivity and coverage for our experimental objectives.
c. *Orbitrap Eclipse gas-phase fractionation (GPF)* was performed using LC-MS settings identical to DIA, except for the following: six narrow MS1 scan ranges (395–505, 495–605, 595–705, 695–805, 795–905, and 895–1005 m/z), precursor isolation windows of 6 Da with 1 Da overlap, 25 MS2 scans per cycle, and a 3-second loop time. The resulting GPF RAW files were used to construct a sample-specific chromatogram library [16] for EncyclopeDIA searches.

### Bioinformatics of peptide spectral matching, protein inference, and quantification

a. *Bioinformatics for DDA MS:* Peptide MS/MS spectra were searched against the UniProt *Homo sapiens* reference proteome (SwissProt, reviewed entries: 20,365 protein sequences, generated on 10 October 2023) appended with reversed decoy and common contaminant (cRAP) database[24] using FragPipe *(v22.0)* [25]. The Raw files were processed using MSFragger (v4.1) for database searching, MSBooster (v1.2.31) for machine-learning score refinement, Percolator (v3.6.5) for rescoring, Philosopher (v5.1.1) for FDR control, ProteinProphet for protein inference, and IonQuant (v1.10.27) for LFQ with match-between-runs (MBR) enabled. Searches used strict trypsin specificity, up to two missed cleavages, precursor tolerance ±20 ppm, fragment tolerance 0.6 Da, fixed carbamidomethylation (+57.021 Da), and variable methionine oxidation (+15.995 Da) and N-terminal acetylation (+42.011 Da) for peptide-spectrum match (PSM) generation and label-free quantification (LFQ), with peptide and protein-level False Discovery Rate (FDR) controlled at 1%. Proteins supported by at least one unique peptide per protein were used for quantification.
b. *Bioinformatics for DIA-MS:* Peptide DIA-MS/MS spectra were searched with EncyclopeDIA (v2.12.30) [16, 26]. To construct a comprehensive chromatogram library, six GPF DIA runs spanning 395–1005 m/z (395–505, 495–605, 595–705, 695–805, 795–905, and 895–1005 m/z) were collectively analyzed as described by Pino et al [26]. Experimental datasets were acquired using wide-window DIA on protein digests from individual BAL fluid and BAL cells. Raw files were converted to the mzML format using msConvert, with peak-picking and denoising filters applied. Predicted spectral libraries (.dlib) were generated from the Human SwissProt proteome database (20,365 protein sequences, generated on 10 October 2023) using the Prosit 2020 Intensity HCD model[27, 28] with parameters allowing up to two missed cleavages, collision energy of 33, peptide lengths between 7 and 30 amino acids, and up to three oxidized methionines. Chromatogram libraries (.elib) were constructed in EncyclopeDIA (v2.12.30) using GPF-derived mzML files, the Prosit-based spectral library, and the SwissProt FASTA database. DIA data from individual samples were searched against the EncyclopeDIA chromatogram library using default tolerances and scoring thresholds, and identifications were filtered to 1% FDR via a target-decoy strategy. The output included peptide- and protein-level quantification tables, sequence information, and fragment ion intensities. Protein quantification within EncyclopeDIA uses at least one peptide per protein, with each peptide having at least three transitions without any fragment ion interference. Peptide quantification within EncyclopeDIA extracts fragment ion chromatograms and applies its built-in interference rejection model to retain only co-eluting, interference-free ions for accurate peak integration [16].

### Statistical analysis

In general, missingness, protein overlap, and abundance distributions were evaluated, with quantification harmonized for comparison. Within BALF, we compared rates of detection/missingness between DIA- and DDA-MS by tabulating the number of proteins that had between zero and five missing values across samples within each method. The proportion of proteins with complete data was compared using chi-squared tests within the BALF and BAL cell sample sets. Proteins with complete data in either DDA- or DIA-MS were categorized into three groups based on whether they were detected by both methods or by only one. For proteins with complete data in both methods, we compared abundance estimates on a log_10_ scale using Bland-Altman analyses. This same procedure was repeated in full using the BAL cells’ protein abundances. Where reported, comparisons of (mean) abundance levels were done via two-sided t-tests of regression coefficients from linear regression models using log_10_-transformed abundance values as the outcomes. To determine the biological relevance of specific sets of proteins (e.g. those detected by DIA- but not by DDA-MS in BALF, etc.), pathway enrichment was performed using Ingenuity Pathway Analysis (IPA) with the core analysis tool, which identifies canonical pathways associated with the selected features.

In accordance with community standards for proteomics data sharing, all raw and processed spectral data have been deposited in the MassIVE repository (MassIVE ID: MSV000100648)

## Results

### BALF peptide and protein identification and quantification across DIA- and DDA-MS

We analyzed spectral data from tryptic digests of medium- and low-abundance BALF proteins using EncyclopeDIA DIA-MS, identifying 1,742 proteins at a 1.0% FDR (Table 1; Supplement 1). The number of proteins quantified across six samples is shown in Figure 1A. Using the same digests, DDA-MS identified 2,069 with FragPipe (Table 1, Supplementary File 1). The FragPipe DDA-MS dataset showed variable protein quantitation, with 62.5% of proteins having no missing values. Specifically, 1,050 proteins were consistently quantified across all six samples, while 202 were missing in at least one sample, 133 in two, 142 in three, and 154 in four samples. In contrast, the DIA-MS dataset analyzed with EncyclopeDIA achieved 97.3% completeness, with 1,695 proteins quantified across all samples. Only 29 proteins were missing in one sample, 6 in two, 7 in three, and 5 in four samples. The difference in proportion of samples with no missing values between DDA- and DIA-MS was 34.8% (95% Confidence Interval: 32.3%-37.3%) with a highly significant chi-squared test (p < 1e-16). Among proteins detected in all samples with DIA- and DDA-MS, 988 were identified by both DIA- and DDA-MS, 707 were unique to DIA-MS, and 62 were exclusive to DDA-MS (Figure 1B). The Bland-Altman analysis for proteins detected in all samples with both DIA- and DDA-MS generally showed strong agreement in quantification (Figure 1C). The mean difference (bias) between methods was close to zero, indicating no systematic shift in protein abundance estimates. The 95% limits of agreement were narrow relative to the data’s dynamic range, indicating strong concordance between the two acquisition strategies.

**Fig 1:**
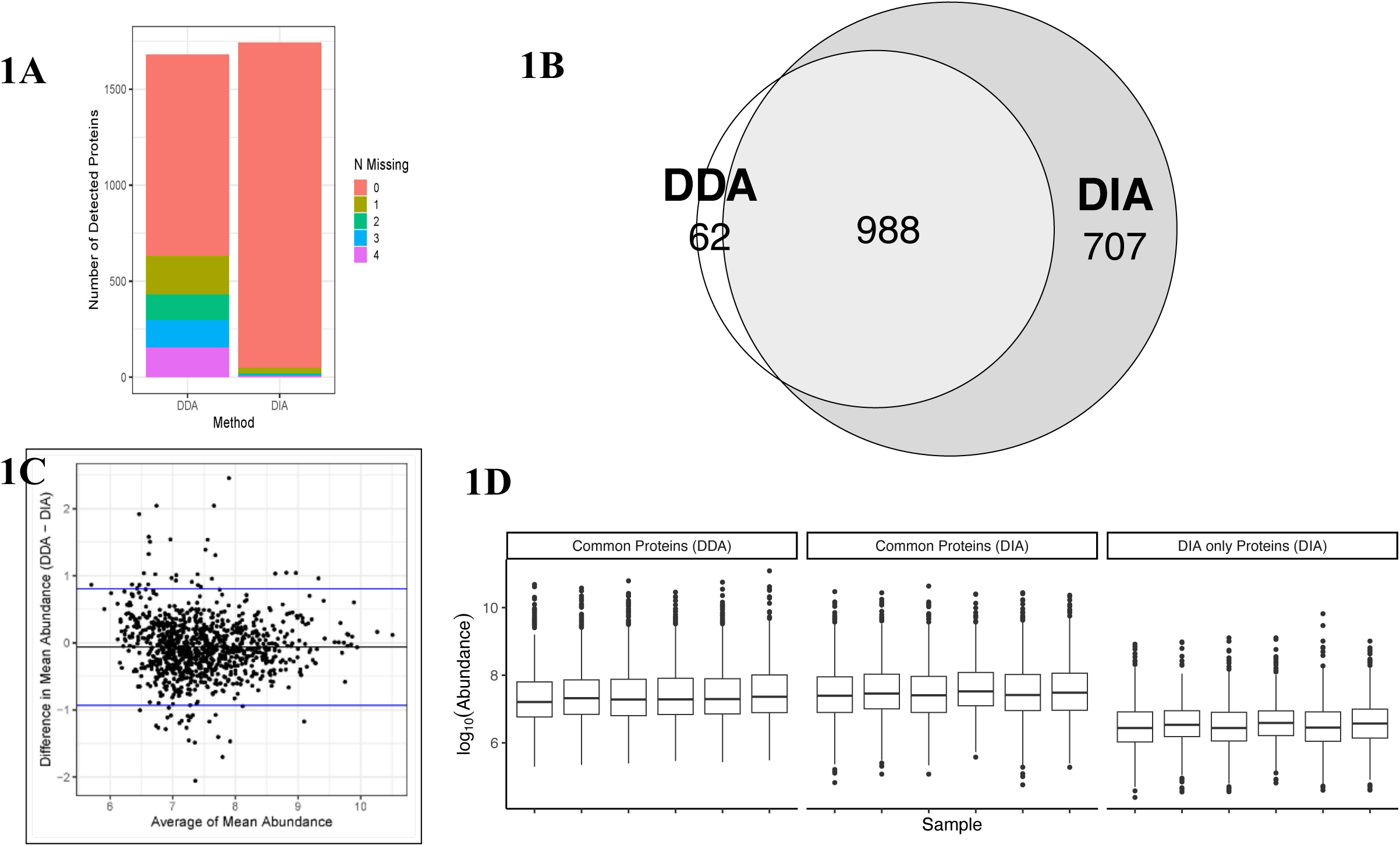
BALF protein coverage using contemporary MS workflows. **Figure 1A**: Number of proteins quantified by EncyclopeDIA DIA-MS and FragPipe DDA-MS, categorized as proteins with no missing values (blue) and proteins with at least one missing value (orange). The absolute number of missing values and the proportion of samples with one to six missing values are also shown. **Figure 1B**: Overlap of proteins that were quantified without any missing values by DIA-MS and DDA-MS. **Figure 1C**: Agreement between DDA and DIA quantification assessed using Bland–Altman analysis. The mean difference (bias) between methods was close to zero, indicating no systematic shift in protein abundance estimates. The 95% limits of agreement were narrow relative to the data’s dynamic range, indicating strong concordance between the two acquisition strategies. Visual inspection of the Bland–Altman plot showed no evidence of proportional bias, and the variance of differences remained stable across the full intensity range. Overall, DDA and DIA produced highly comparable quantitative values for proteins detected by both workflows. **Figure 1D**: The abundance of proteins that were identified by both FragPipe DDA and EncyclopeDIA, quantified by either method, was higher than the abundance of proteins uniquely detected by EncyclopeDIA. Each sub-panel describes which proteins are being shown, as well as which quantification method is being used in parentheses.

**Table 1:** BALF proteins identified and quantified by various PSM search algorithms for DDA- and DIA-MS.

| TOOLS | Analysis | Peptides | Detected Proteins | Fully Quantified Proteins | Missing Values Proteins | No Quant Values Proteins | Percent missing |
| --- | --- | --- | --- | --- | --- | --- | --- |
| EncyclopeDIA | DIA | 12432 | 1742 | 1695 | 47 | 0 | 2.7% |
| FragPipe | DDA | 16629 | 2069 | 1050 | 631 | 388 | 37.5% |

*Detected – identified in output*
*Fully quantified – all six samples have intensities*
*Missing values – protein has at least one missing value, but not all six*
*No quant – no intensities are listed for that protein*
*Percent missing – based on proteins that have quantification, did not include proteins with no quant*

Visual inspection of the Bland–Altman plot showed no evidence of proportional bias, and the variance of the differences remained stable across the full intensity range. Overall, DDA- and DIA-MS produced highly comparable quantitative values for proteins detected by both workflows. However, proteins uniquely detected by DIA-MS tended to be less abundant (Figure 1D) than those that were detected by both DIA- and DDA-MS. The mean abundance levels, as quantified using DIA-MS, for the proteins unique to DIA-MS were lower by 0.97 units (log_10_ scale) on average than those that were common to DIA- and DDA-MS (p < 1e-16).

### Comparison of BAL cell proteins identified and quantified by DIA-MS and DDA-MS

We identified 5,227 high-confidence proteins (FDR < 1.0%) using EncyclopeDIA, while FragPipe detected 5,640 proteins. DIA-MS provided quantitative data for all 5,227 identified proteins, with only 49 proteins (0.94%) missing, resulting in 5,178 proteins (99.06%) quantified across all samples. Missing values were distributed across 25 proteins in one sample, 9 in two samples, 5 in three samples, 7 in four samples, and 3 in five samples. In contrast, DDA-MS yielded quantitative data for 4,894 of the 5,640 identified proteins (Supplementary File 2, Table 2). FragPipe had missing values for 1,355 proteins (27.7%), leaving 3,539 proteins (72.3%) fully quantified across all samples (Figure 2A). Missing values were observed for 448 proteins in one sample, 360 in two samples, 266 in three samples, and 281 in four samples, with no proteins missing across five samples. The difference in the proportion of proteins with no missing values between DDA-MS and DIA-MS was 26.7% (95% confidence interval: 25.4–28.0%), with a highly significant chi-squared test (p < 1 × 10⁻¹⁶). Among proteins fully quantified across all samples, 3,397 were shared between DIA-MS and DDA-MS. Additionally, 1,781 proteins were uniquely detected by DIA-MS, while 142 proteins were uniquely detected by DDA-MS (Figure 2B). The Bland-Altman analysis of proteins detected by both DIA- and DDA-MS with complete data was virtually identical to what was observed in BALF (Figure 2C). The bias was again almost zero, the 95% agreement limits were very similar, and neither the bias nor the variance seem to depend on the mean abundance value. Also, as with BALF, the BAL cell proteins detected only by DIA-MS were lower in abundance than those detected by DDA-MS (Figure 2D). The mean abundance levels, as quantified using DIA-MS, for the proteins unique to DIA-MS were lower by 0.89 units (log_10_ scale) on average than those that were common to DIA- and DDA-MS (p < 1e-16).

**Fig 2:**
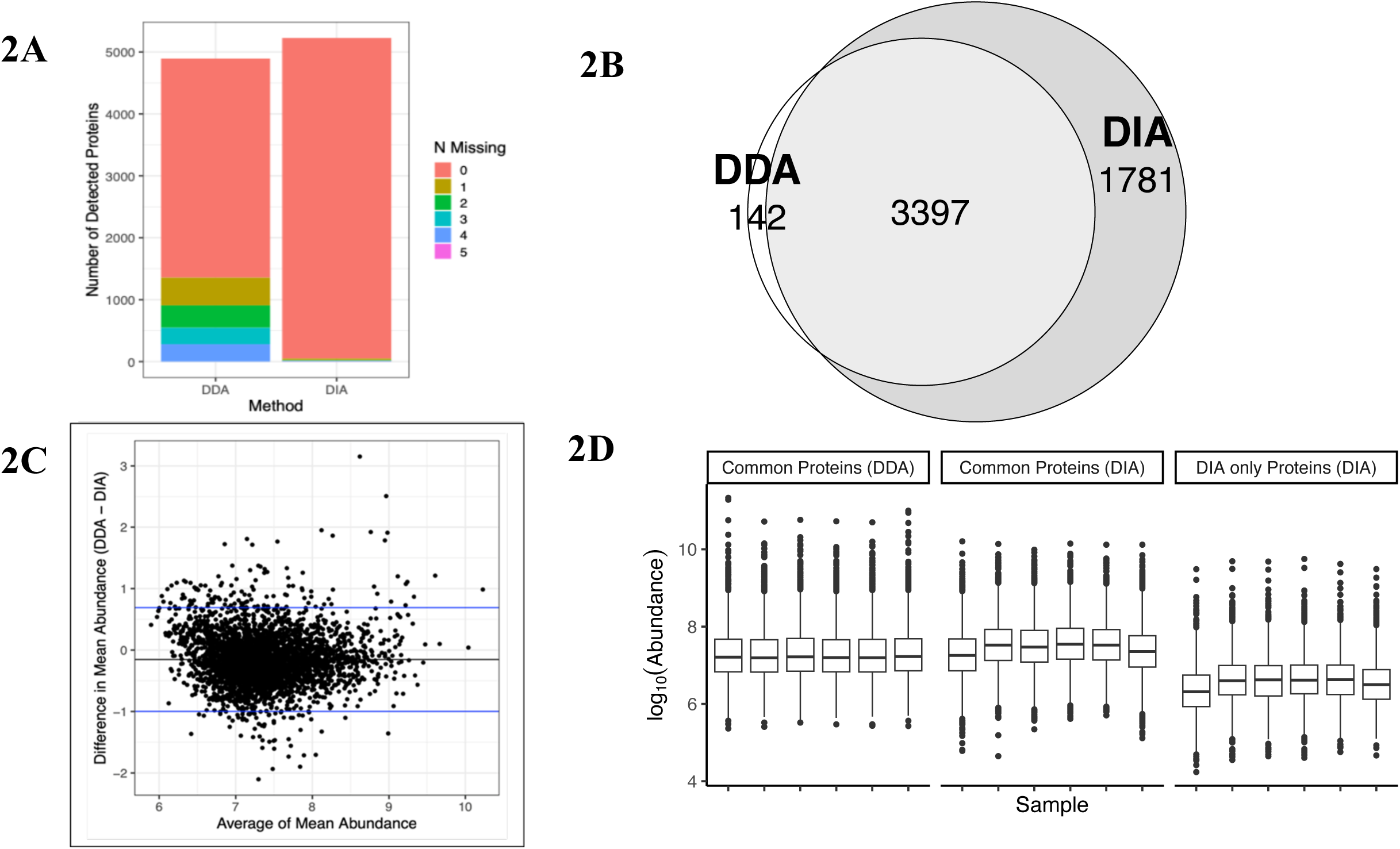
BAL cell protein coverage using contemporary MS workflows. **Fig 2A** Number of proteins quantified by Encyclopedia DIA-MS and FragPipe DDA-MS, categorized as proteins with no missing values (blue) and proteins with at least one missing value (orange). The absolute number of missing values and the proportion of samples with one to six missing values are also shown. **Figure 2B**: Overlap of proteins that were quantified without any missing values by DIA-MS and DDA-MS. **Figure 2C**: Agreement between DDA and DIA quantification assessed using Bland–Altman analysis. The mean difference (bias) between methods was close to zero, indicating no systematic shift in protein abundance estimates. The 95% limits of agreement were narrow relative to the data’s dynamic range, indicating strong concordance between the two acquisition strategies. Visual inspection of the Bland–Altman plot showed no evidence of proportional bias, with stable variance across intensity range. Overall, DDA and DIA produced highly comparable quantitative values for proteins detected in both workflows. Figure 2D: **Figure 2D**: The abundance of proteins quantified by FragPipe DDA, either uniquely or in conjunction with EncyclopeDIA, was higher than the abundance of proteins uniquely detected by EncyclopeDIA.

**Table 2:** BAL cell proteins identified and quantified by various PSM search algorithms for DDA- and DIA-MS.

| TOOLS | Analysis | Peptides | Detected Proteins | Fully Quantified Proteins | Missing Values Proteins | No Quant Values Proteins | Percent missing |
| --- | --- | --- | --- | --- | --- | --- | --- |
| EncyclopeDIA | DIA | 42650 | 5227 | 5178 | 49 | 0 | 0.94% |
| FragPipe | DDA | 48922 | 5640 | 3539 | 1355 | 746 | 27.7% |

*Detected – identified in output*
*Fully quantified – all six samples have intensities*
*Missing values – protein has at least one missing value, but not all six*
*No quant – no intensities are listed for that protein*
*Percent missing – based on proteins that have quantification, did not include proteins with no quant*

We analyzed proteins from EncyclopeDIA and FragPipe without missing values using IPA Core Analysis. Canonical pathway enrichment revealed that proteins identified in BAL cells mapped to biological pathways previously implicated in granulomatous inflammatory processes. We then performed a comparison analysis between DIA- and DDA-based proteomes to gain insights into biological processes captured by the two MS acquisition techniques (Supplementary File 3). Shared pathways between proteins identified by DIA- and DDA-MS included EIF2 signaling, TLR signaling, clathrin-mediated endocytosis, sirtuin signaling, mTOR signaling, C-type lectin receptor signaling, and others (Supplemental Table 3, worksheet ‘Significant in Both’). Many pathways in the overrepresentation analysis were statistically significant for DIA-MS proteins compared with those identified by DDA-MS (Table 3). In contrast, some pathways were significant in DDA-MS but not in DIA-MS (Supplemental File 3, spreadsheet ‘Significant in DDA Only’). We then performed IPA Core Analysis using the 1,781 proteins identified exclusively by DIA-MS. Pathways mapped to these DIA-only proteins included the complement cascade, regulation of TLR signaling by endogenous ligands, the coagulation system, and JAK/IL-6-type cytokine signaling (Figure 3, Supplementary Table 3, spreadsheet ‘Pathways DIA-only proteins’)

**Fig 3:**
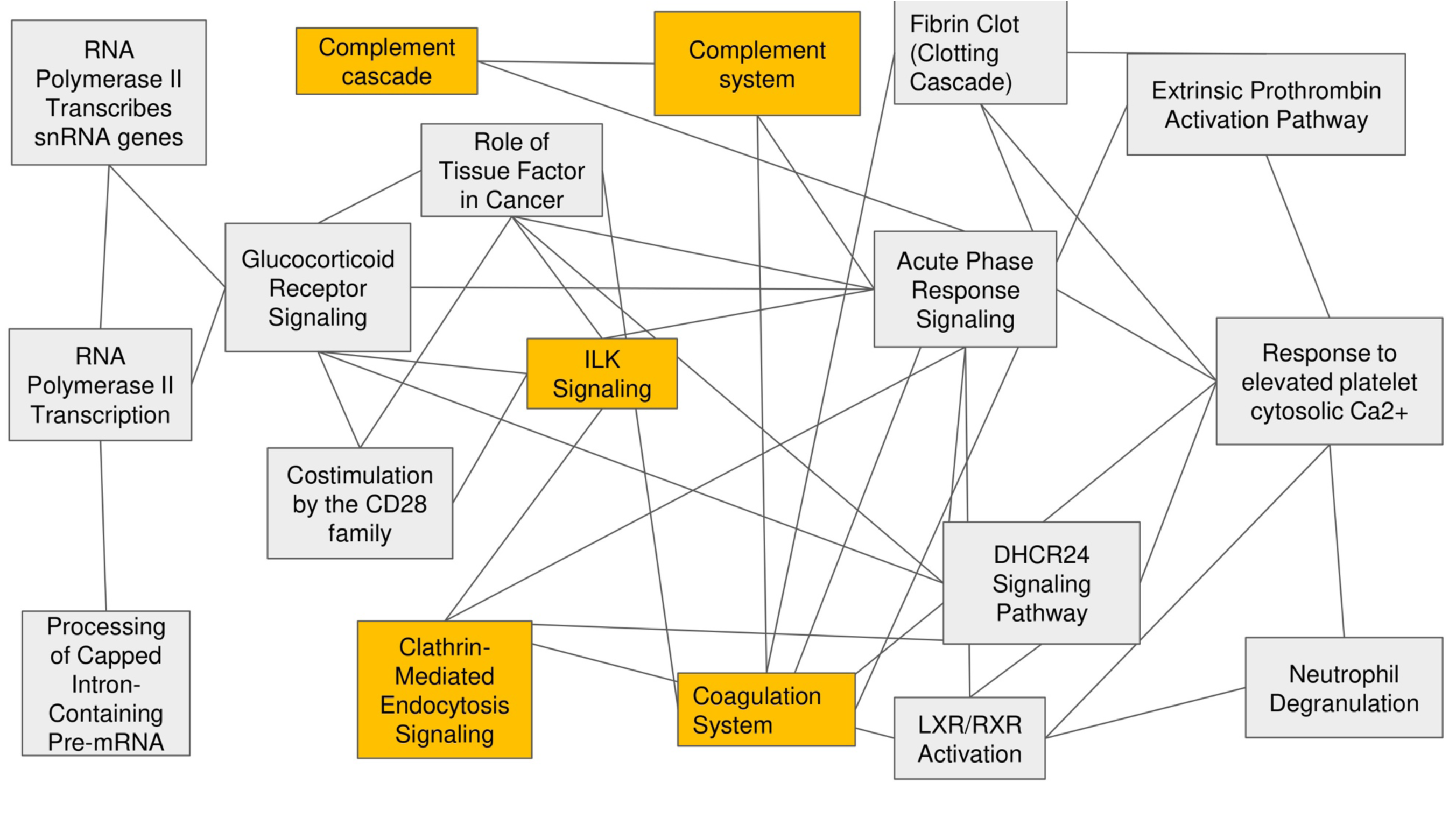
Ingenuity Pathway Analysis (IPA) Core Analysis of BAL cells, visualized using the Overlapping Pathways tool, identified canonical pathways enriched exclusively through DIA-MS. DIA-MS uniquely quantified 1,781 proteins, enabling a more comprehensive interrogation of pathway-level biology compared to data-dependent methods. Shared canonical pathways included EIF2 signaling, Toll-like receptor (TLR) signaling, clathrin-mediated endocytosis, sirtuin signaling, and C-type lectin receptor (CLR) signaling. In contrast, DIA-MS uniquely revealed enrichment of pathways such as the complement cascade, regulation of TLR signaling by endogenous ligands, the coagulation system, and the role of Janus kinase (JAK) kinases in interleukin-6 (IL-6)-type cytokine signaling. In the graphical representation, each node denotes a canonical pathway, whereas connecting edges reflect curated functional or regulatory relationships between pathways derived from IPA’s integrated molecular

**Table 3:**
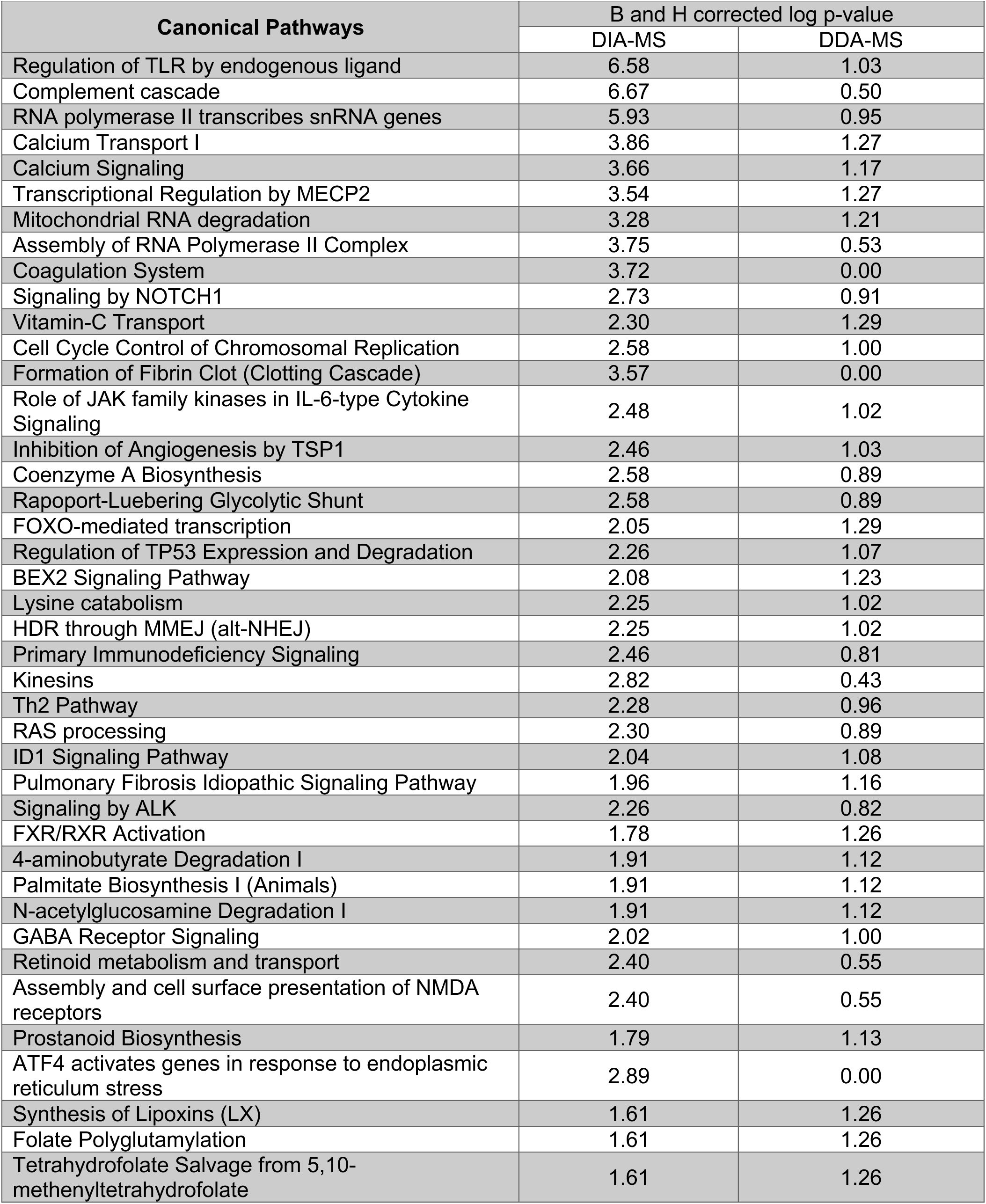
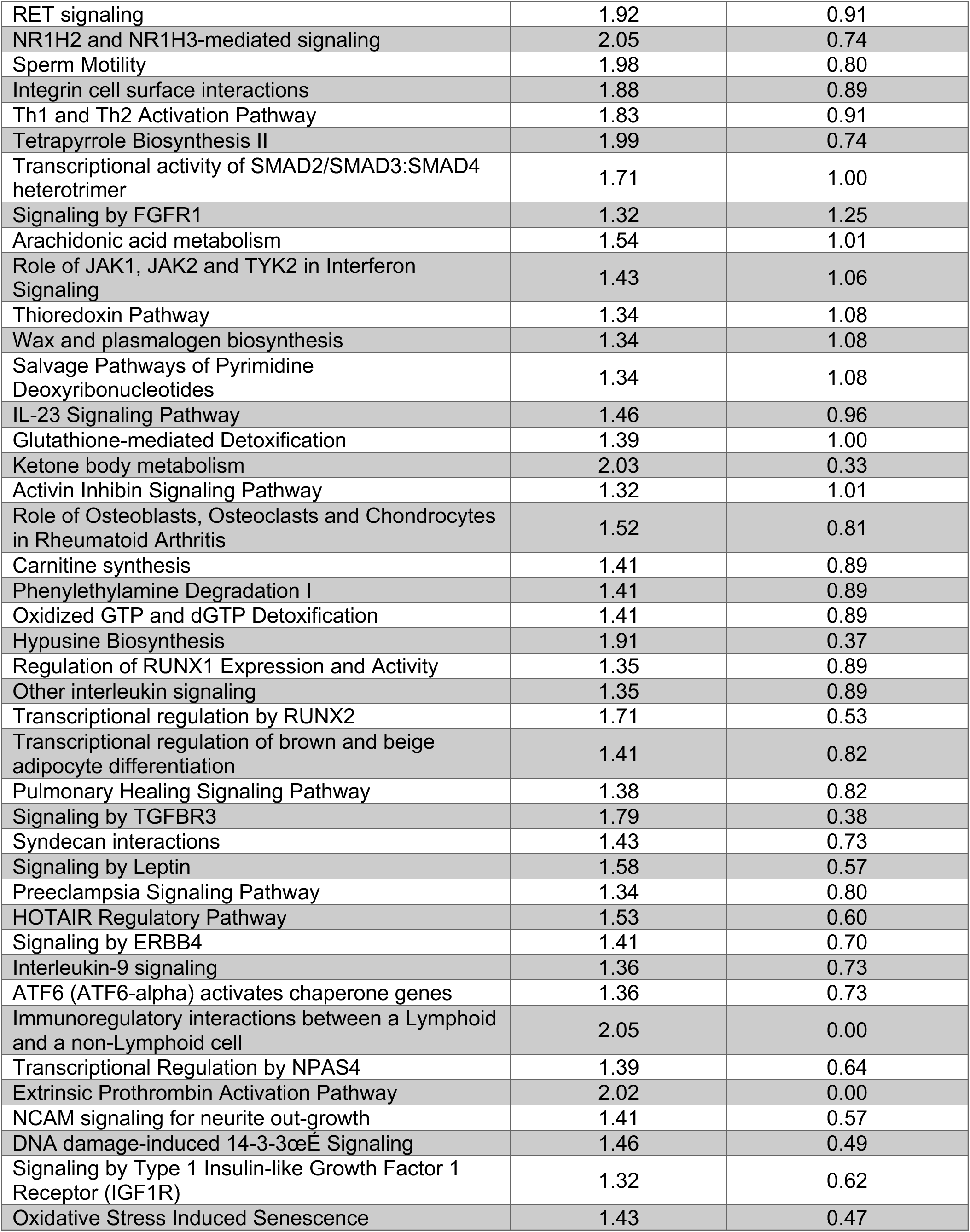
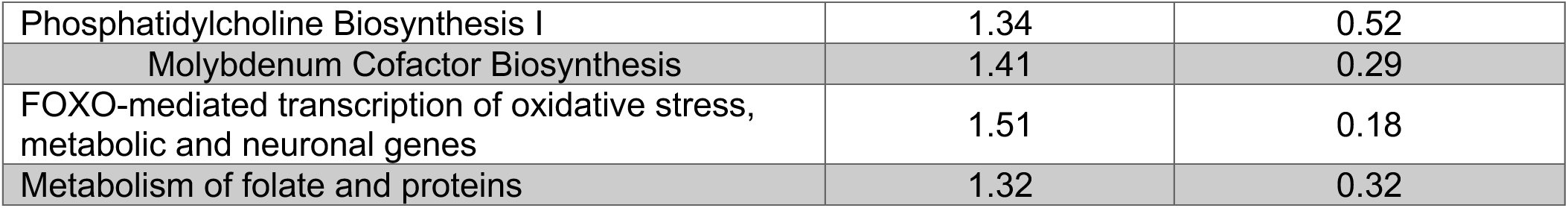
Canonical Pathways mappting to proteins detected by both DIA- and and DDA-MS but do not reach significance threshold for proteins identified by.

### Impact of duration and method of biospecimen storage on protein yield

The number of proteins quantified with EncyclopeDIA in BALF samples from 2003, 2012, and 2023 was remarkably consistent at 1,736, 1,738, and 1,736, respectively. Additionally, for the paired BAL cells, the numbers of detected proteins were 5,216, 5,219, and 5,225. When comparing the cryopreserved BAL cells with flash-frozen cells, the number of proteins identified in the cryopreserved cells was 2,478, and in flash-frozen cells was 2,403, with 2,283 (87.8 %) of the proteins detected, with cryopreserved samples having 195 unique proteins and flash-frozen samples having 121 unique proteins. (Figure 4, Supplementary Table 4)

**Fig 4:**
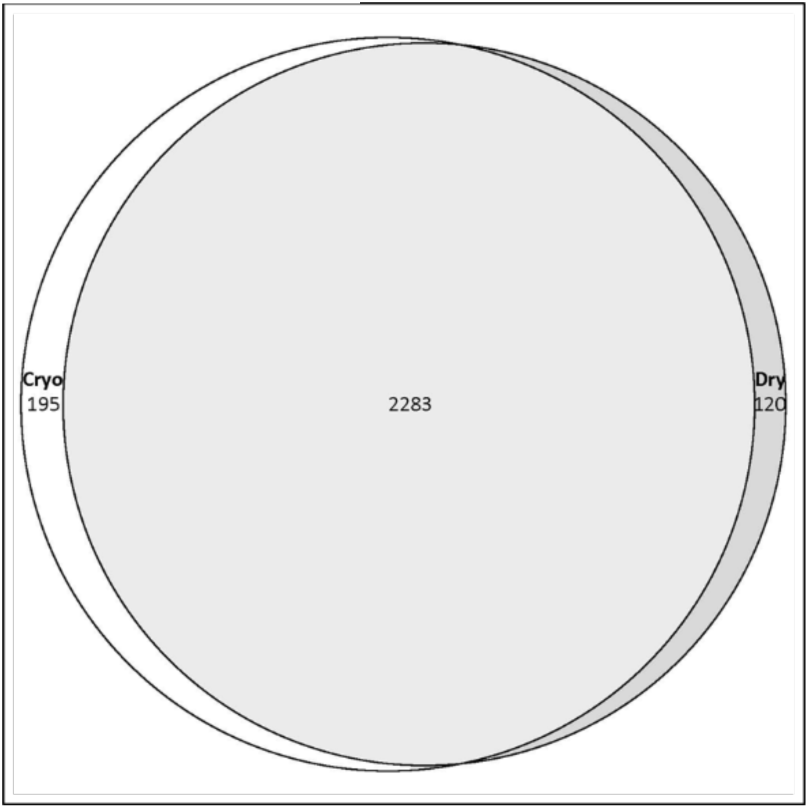
BAL cells from three samples stored through cryopreservation or rapid freezing in liquid nitrogen were compared for protein yield. The majority of the proteins (87.8%) were detected in both storage methods, with only 195 proteins uniquely identified in cryopreserved cells and 120 proteins found in flash-frozen cell

### Comparing paired BAL cell and BALF samples

The total number of proteins fully quantified by DIA-MS was higher in BAL cells (5,178) than in BALF (1,695). Although the DIA-MS detected 1,234 proteins in both BAL cells and BALF, 461 proteins were unique to BALF, and 3,944 proteins were exclusive to BAL cells (Figure 5, Supplemental Table 5).

**Figure 5:**
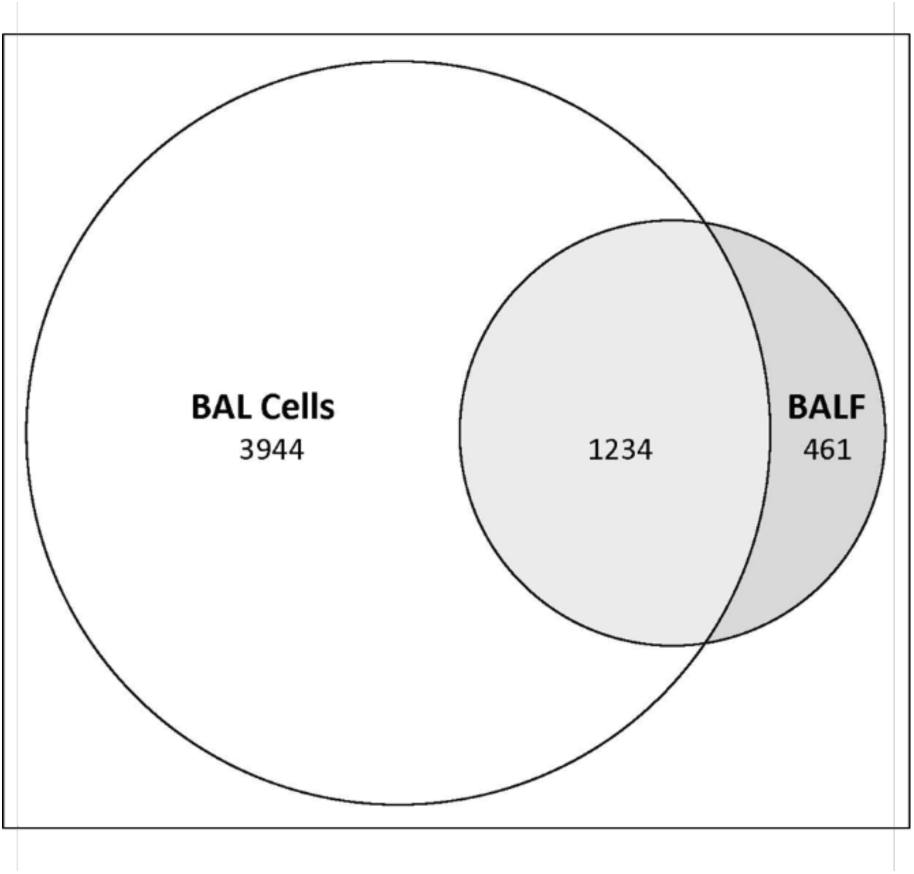
Paired BAL cells and BALF samples were examined. The number of proteins detected by EncyclopeDIA in BAL cells was 5,178, and in BALF was 1,695. Although some unique proteins were identified in BALF, most of the proteins found in BALF were also present in BAL cells, with many proteins detected only in BAL cells.

## DISCUSSION

Significant progress has been made in recent years in characterizing complex protein mixtures, driven by advances in MS technologies. Modern workflows combine comprehensive MS acquisition strategies, such as DIA-MS, with sophisticated algorithms for peptide spectral matching and protein inference. These developments have improved proteome coverage, quantitative accuracy, and reproducibility, promising deeper insights into biological systems and disease mechanisms. BALF is inherently difficult to analyze by mass spectrometry because of its wide dynamic range, in which high-abundance proteins and complex fluid components obscure detection of lower-abundance proteins and limit proteome depth. To guide the selection of acquisition strategies for robust characterization of lung compartment changes, we systematically analyzed BALF and BAL cells to inform ongoing studies. Although we found protein identification was similar between DDA and DIA-MS, quantification was more robust with DIA-MS, with fewer missing values. In both BAL cells (Fig 1D) and BALF (Fig 2D), many lower-abundance proteins were detected only by DIA-MS. While DIA-MS involves more computationally intensive data processing than traditional DDA-MS, advances in algorithms and library-free approaches are reducing these barriers [29]. As workflows mature, DIA’s advantages in reproducibility, depth, and pathway resolution make it well-suited for translational studies in lung immunology and beyond.

The challenges and value of BALF for MS-based proteomics have been well recognized [30, 31]. These challenges arise from the chemical complexity of epithelial lining fluid and include high-abundance proteins that obscure detection of lower-abundance lung proteins; interfering substances such as mucus, lipid surfactants, and high salt from saline used during collection, all incompatible with nanoscale LC–MS/MS systems; dilution of tissue-derived molecules due to large saline volumes; and limited protein yield from specific subjects or disease states, restricting deep proteome coverage. Although BALF is the most proximal fluid at the site of lung damage and provides unique insights into local pathophysiology, its collection requires invasive bronchoscopy. Therefore, leveraging archived biospecimens collected over many years is advantageous to enable robust analyses without additional patient risk and to ensure adequate statistical power for patient-oriented studies. Our group previously reported an optimized workflow for BALF processing[19]. The high-quality peptide mixtures generated by this workflow are amenable to either semi-preparative offline high-pH HPLC fractionation (for samples yielding tens of micrograms of peptide) or micro-fractionation (for samples yielding as little as 1 μg of peptide). As a logical next step, we compared DIA-MS and DDA-MS performance and observed similar protein identification from trypsin-digested peptides across medium- and low-abundance proteins, with the use of EncyclopeDIA and FragPipe, both of which are open source, and EncyclopeDIA is a foundational tool optimized for DIA-MS analysis, while FragPipe is a leading tool for DDA-MS[17, 32]. However, DIA-MS offered deeper coverage and fewer missing values than DDA-MS, similar to prior observations in other tissue matrices [15, 33]. Many of the additional proteins detected by DIA-MS were less abundant than those identified by DDA-MS, highlighting the advantage of DIA in capturing proteins that might otherwise be missed in traditional acquisition workflows. Such workflows can be scaled to large sample sizes for MS-based proteomics studies aimed at biomarker discovery. A key advantage of label-free quantification approaches, such as DIA-MS, is their ability to generate peptide transition information, which supports analyte-level validation for targeted proteomics, and facilitates the development of clinically translatable assays, including those that are suitable for clinical validation and use [34, 35].

Assessment of BAL cell protein is particularly important because it provides direct information about inflammatory processes within the lung. Most BAL cells are leukocytes, including alveolar macrophages, lymphocytes, and neutrophils, and reflect immune activation and inflammatory processes within the lung microenvironment. Therefore, proteomic analysis of BAL cells offers unique insights into intracellular signaling, immune regulation, and cellular stress responses that are not captured by BALF alone. However, as with BALF, BAL cell proteomics faces challenges, including limited sample volumes and the need for invasive procedures for biospecimens procurement. In our study, DIA-MS demonstrated superior performance in BAL cell analysis compared to DDA-MS, offering deeper coverage and fewer missing values, particularly for low-abundance proteins involved in immune and metabolic pathways. Moreover, DIA-MS provided broader coverage of canonical pathways than DDA-MS, enabling a more comprehensive view of the intricate protein interaction network. This enhanced pathway resolution offers deeper insights into immune and metabolic processes that may underlie granulomatous inflammation and may be implicated in beryllium-related health effects. Specifically, DIA-MS captured nearly 1,781 additional proteins—primarily low-abundance species—while maintaining high overlap with DDA. Improved quantitation, resulting from reduced missingness and greater sensitivity, translated into enhanced biological resolution, particularly in immunologically relevant pathways. Our observed comparability between cryopreserved and flash-frozen samples validates use of existing repositories for retrospective studies, expanding research opportunities.

We also evaluated the duration of sample storage via cryopreservation or flash-freezing, with no significant difference, indicating that both preservation methods are suitable for downstream LC-MS analysis of inflammatory cell populations in the lung. Thus, BAL cells collected previously can be considered for examination with contemporary MS-proteomics tools, regardless of the method of storage. These analyses were performed to assess technical robustness and were not used for biological inference.

## CONCLUSION

DIA-MS provides robust, sensitive, and biologically informative proteomic profiling of BAL cells and fluid in BeS. Compared with DDA, DIA identifies low-abundance proteins, reduces missing data, and enhances the resolution of key inflammatory pathways. These strengths position DIA-MS as a well-suited platform for biomarker discovery and mechanistic insights into antigen-mediated granulomatous lung disease. Furthermore, these findings suggest that previously frozen samples perform well, supporting their use in studies that reduce the need for additional procedures, and that cryopreservation may not confer any benefit, thereby minimizing processing time. Future integration of DIA with multi-omics will further clarify disease pathogenesis and reveal therapeutic targets.

## SUPPORTING INFORMATION

List of supplemental material

1. Supplemental Doc: Additional methods and results
2. Supplementary File 1: DIA and DDA protein level data for BALF obtained from multiple analytical platforms.
3. Supplementary File 2: DIA and DDA protein level data for BAL cells obtained from multiple analytical platforms.
4. Supplementary File 3: IPA Canonical Pathways mapping to proteins. Comparison analysis between DIA- and DDA- based proteins.
5. Supplementary File 4: DDA unique and shared protein level data for BAL cell cryo and dry pellets.
6. Supplementary File 5: EncyclopeDIA unique and shared protein level data for BALF and BAL cells.

## NOTES

### Other contributions

The Orbitrap Eclipse instrumentation platform used in this work was purchased through High-end Instrumentation Grant S10OD028717 from the NIH. The studies were conducted in Center of Mass Spectrometry and Proteomics at the University of Minnesota, and we acknowledge the staff for their support in the execution of these studies.

### CRediT Authorship Contribution Statement

**DOW** ORCID: 0000-0001-8644-8030; Investigation (proteomics sample processing), Data Curation, Validation, Visualization, Writing – Original Draft, Writing – Review & Editing

**KG** ORCID ID: 0000-0002-2099-6306; Visualization, Writing – Original Draft, Writing – Review & Editing

**TJG** ORCID: 0000-0001-6801-2559; Methodology, Formal Analysis, Data Curation, Visualization, Writing – Review & Editing

**PDJ** ORCID:0000-0003-0984-0973; Methodology, Formal Analysis, Data Curation, Visualization, Writing – Review & Editing

**MMM** ORCID: 0000-0001-5252-5990; Resources, Data Curation, Writing – Review & Editing

**RW** ORCID: 0009-0008-9774-9291; Data Curation, Software, Writing – Review & Editing

**JDM** ORCID:0000-0003-0664-1178; Resources, Writing – Review & Editing

**SM** ORCID: 0000-0001-9818-0537; Methodology, Formal Analysis, Data Curation, Visualization, Writing – Review & Editing

**LAM** ORCID: 0000-0001-6872-1769; Funding Acquisition, Sample acquisition, Resources, Writing – Review & Editing

**LL** ORCID: 0000-0002-0692-6889; Funding Acquisition, Project Administration, Resources, Writing – Review & Editing

**BV** ORCID: 0000-0002-3772-1691; Funding Acquisition, Methodology, Formal Analysis, Validation, Visualization, Writing – Review & Editing

**MB** ORCID: 0000-0002-1294-6181; Conceptualization, Funding Acquisition, Supervision, Methodology, Validation, Resources, Project Administration, Writing – Original Draft, Writing – Review & Editing

## Supporting information

Supplemental Doc: Additional methods and results

