## Supplemental Doc: Additional methods and results for "Comparative Evaluation of DDA- and DIA-Based Proteomic Workflows in Beryllium-Related Lung Disease"

a. DDA Bioinformatics: For peptide spectral matching of DDA-MS data, in addition to FragPpie DDA, we conducted database search using Proteome Discoverer (v3.1.1.93): Data were searched using SEQUEST with precursor recalibration. Parameters included trypsin specificity, up to two missed cleavages, precursor tolerance 15 ppm, fragment tolerance 0.6 Da, fixed carbamidomethylation (+57.021 Da), and dynamic modifications (N-terminal acetylation, methionine oxidation, pyroglutamate formation, methionine loss ± acetylation, and deamidation). LFQ was performed using Minora Feature Detector for feature extraction, chromatographic alignment, normalization, and protein abundance calculation.

b. Bioinformatics for DIA: We also conducted peptide spectral matching for the DIA-MS data using *PD CHIMERYS* 3.0.0[1]*:* Peptide DIA MS/MS data were processed in Proteome Discoverer (PD v3.1.1.93, Thermo Scientific) using the CHIMERYS spectral prediction algorithm (v3.0.0; https://www.msaid.de/). Searches were performed against the UniProt human reference proteome (UP000005640, downloaded on August 18, 2023), merged with a common contaminant databas[2 3] . CHIMERYS parameters included the inferys_3.0.0 prediction model, trypsin specificity with up to one missed cleavage, peptide lengths of 7–30 amino acids, precursor charge states 1–4, and fragment ion mass tolerance of 10 ppm. Carbamidomethylation of cysteine (+57.021 Da) was specified as a fixed modification, while oxidation of methionine (+15.995 Da) was set as a variable modification (maximum three per peptide). LFQ was performed in PD using fragment ion intensities of unique and razor peptides, excluding shared peptides. Normalization was applied to summed peptide fragment ion abundances, and protein ratios were calculated using a pairwise ratio-based approach similar to MaxLFQ[4] , controlling for FDR.

**Additional Results**

BALF protein identification and quantification: Spectral data from tryptic digests of medium- and low-abundance BALF proteins were analyzed using DIA-MS. EncyclopeDIA identified 1,742 proteins and PD Chimerys identified 1,536 proteins at an FDR of 1.0% (Table 1S; Supplement 1). EncyclopeDIA quantified 1,695 proteins with no missing values (2.7% missingness), whereas PD Chimerys quantified 1,180 proteins without missing values, corresponding to 23.3% missingness. Using DDA-MS on the same tryptic digested peptides, FragPipe identified 2,069 proteins and PD identified 1,938 proteins at an FDR of 1%. In FragPipe DDA, 1,050 proteins were fully quantified, with no quantitative data for 388 proteins and at least one missing value in 631 proteins. For PD, 510 proteins lacked quantitative data, and 804 had at least one missing value.

**Table 1S: BALF proteins identified and quantified by various PSM search algorithms for DDA- and DIA-MS**

| TOOLS | Analysis | Peptides | Detected Proteins | Fully Quantified Proteins | Missing Values Proteins | No Quant Values Proteins | Percent missing |
|---|---|---|---|---|---|---|---|
| EncyclopeDIA | **DIA-MS** | 12432 | 1742 | 1695 | 47 | 0 | 2.7% |
| PD Chimerys | | 14475 | 1536 | 1180 | 356 | 0 | 23.2% |
| Fragpipe DDA | **DDA-MS** | 16629 | 2069 | 1050 | 631 | 388 | 37.5% |
| PD DDA | | 18733 | 1938 | 624 | 804 | 510 | 56.3% |

*Detected – identified in output*
*Fully quantified – all six samples have intensities*
*Missing values – protein has at least one missing value, but not all six*
*No quant – no intensities are listed for that protein*
*Percent missing – based on proteins that have quantification, did not include proteins with no quant*
